# Enterococcus faecalis antagonizes Pseudomonas aeruginosa growth in polymicrobial biofilms

**DOI:** 10.1101/2022.01.18.476859

**Authors:** Casandra Ai Zhu Tan, Ling Ning Lam, Goran Biukovic, Eliza Ye-Chen Soh, Xiao Wei Toh, José A. Lemos, Kimberly A. Kline

## Abstract

*Enterococcus faecalis* is often co-isolated with *Pseudomonas aeruginosa* in polymicrobial biofilm-associated infections of wounds and the urinary tract. As a defense strategy, the host innately restricts iron availability at infection sites. Despite their co-prevalence, the polymicrobial interactions of these two pathogens in iron-restricted conditions, such as those found in the host, remains unexplored. Here we show that *E. faecalis* inhibits *P. aeruginosa* growth within biofilms when iron is restricted. *E. faecalis* lactate dehydrogenase (*ldh1*) gives rise to L-lactate production during fermentative growth. We find that *E. faecalis ldh1* mutant fails to inhibit *P. aeruginosa* growth. Additionally, we demonstrate that *ldh1* expression is induced in iron-restricted conditions, resulting in increased lactic acid exported and consequently, a reduction in pH. Together, our results suggest that *E. faecalis* synergistically inhibit *P. aeruginosa* growth by decreasing environmental pH and L-lactate-mediated iron chelation. Overall, this study highlights that the microenvironment in which the infection occurs is important for understanding its pathophysiology.

**IMPORTANCE:** Many infections are polymicrobial and biofilm-associated in nature. Iron is essential for many metabolic processes and plays an important role in controlling infections, where the host restricts iron as a defense mechanism against invading pathogens. However, polymicrobial interactions between pathogens are underexplored in iron-restricted conditions. Here, we explore the polymicrobial interactions between commonly co-isolated *E. faecalis* and *P. aeruginosa* within biofilms. We find that *E. faecalis* modulates the microenvironment by exporting lactic acid which further chelates already limited iron, and also lowers the environmental pH to antagonize *P. aeruginosa* growth in iron-restricted conditions. Our findings provide insights into polymicrobial interactions between pathogens in an infection-relevant condition and how manipulating the microenvironment can be taken advantage of to better control infections.

## INTRODUCTION

Many infections are often polymicrobial in nature (1–3) and include wound infections (4–7), periodontitis (8, 9), otitis media (10, 11), urinary tract infections (UTI) (12–15) and cystic fibrosis (16–20). Biofilms are also implicated in all these infections (21–37). Polymicrobial biofilms can better tolerate antibiotic treatment and escape from host immune responses, enabling the survival and persistence of the infecting bacteria (38, 39). Understanding how pathogens interact during infections may inform improved treatment strategies.

Iron is an essential nutrient for almost all microbial species. In humans, iron regulation functions as a host innate immune mechanism against invading pathogens (40). In the human body, iron is scarcely available to pathogens due to the sequestration of most iron intracellularly such that only a small amount of free iron (approximately 10^-24^ M) is accessible in the absence of infection (41). During an infection, additional iron-withholding mechanisms further restrict iron availability to pathogens (40). For example, immune cells producing hepcidin (42) or lactoferrin (43), and siderocalin/lipocalin-2 (43, 44) modulate iron availability at the infection site. As a result, when developing polymicrobial interaction models, it is critical to take into account the iron availability in the environment.

Enterococci are opportunistic pathogens implicated in several types of infections (45), and enterococcal infections in humans are mostly caused by *Enterococcus faecalis* and *Enterococcus faecium* (46, 47). *E. faecalis* is often isolated from infective endocarditis, UTI and mixed-species chronic wounds (46, 48, 49). In addition to *E. faecalis,* infected wounds often contain other bacteria such as *P. aeruginosa, S. aureus, Corynebacterium spp, Enterobacteriaceae spp*, and *Finegoldia magna* (4, 48, 50–52). *E. faecalis* co-infection with *S. aureus*, *P. aeruginosa* and *F. magna* resulted in delayed wound closure and higher level of antimicrobial tolerance (52). During UTI, *E. faecalis* is also co-isolated with *P. aeruginosa* and *Proteus mirabilis* (53, 54). *E. faecalis* and *P. aeruginosa* co-infection during UTI resulted in more severe lesions in the kidneys and *P. aeruginosa* was more resistant to clearance by β-lactam antibiotics (53). While increasing efforts have been made to understand how interactions between pathogens can influence bacterial pathogenesis, few studies have explored the polymicrobial interactions in an infection-relevant, iron-restricted condition. Since *E. faecalis* and *P. aeruginosa* are commonly co-isolated in different infection niches, understanding the polymicrobial interactions between these two species is of interest and clinical relevance. Further, there have been no reports examining the interaction between these two species in iron-restricted conditions in any infection model.

In this work, we showed that *E. faecalis* inhibited *P. aeruginosa* growth within biofilms when iron was restricted. The growth inhibition was a consequence of increased L-lactate produced by *E. faecalis*, catalysed by lactate dehydrogenase (*ldh1*) from pyruvate. We also showed that *ldh1* expression was upregulated when iron was restricted. L-lactate produced by *E. faecalis* was exported from the cell as lactic acid (55), whereupon it was deprotonated to L-lactate releasing a hydrogen ion (H^+^) and in turn lowering the pH in the surrounding environment. We demonstrated that this lowered environmental pH and L-lactate-mediated chelation of iron ultimately contributes to *P. aeruginosa* growth inhibition by *E. faecalis* in iron-restricted conditions.

## RESULTS

### *E. faecalis* inhibits *P. aeruginosa* growth in iron-restricted conditions

The compound 2,2’-dipyridyl (22D) is widely used as a neutral ligand for the chelation of metal ions and is a high affinity chelator of iron (56). Therefore, 22D was supplemented into TSBG growth media to restrict iron availability. We first investigated the interactions between 12 different *E. faecalis* clinical isolates and *P. aeruginosa* PAO1 BAA-47 using static biofilm assays. In iron-restricted media (TSBG supplemented with 1 mM 22D), PAO1 BAA-47 growth was inhibited in all of the mixed-species biofilms compared to PAO1 BAA-47 single-species biofilm (**Figure 1A**). By contrast, *E. faecalis* growth was similar in the single- and mixed-species biofilms (**Figure 1B**). These data indicate that *E. faecalis* inhibition of *P. aeruginosa* is not strain specific. We next performed the biofilm assay, now with *E. faecalis* OG1RF and eight different *P. aeruginosa* clinical isolates, to examine whether *P. aeruginosa* susceptibility to *E. faecalis*-mediated inhibition was strain specific. We observed that the growth of all *P. aeruginosa* isolates was inhibited in the mixed-species biofilms compared to their respective single-species counterpart (**Figure 1C**), while *E. faecalis* growth in the single- and mixed-species biofilms remained unaffected (**Figure 1D**). These data demonstrate that all tested *E. faecalis* and *P. aeruginosa* clinical isolates engage in mixed-species antagonism in iron-restricted conditions.

**Figure 1.**
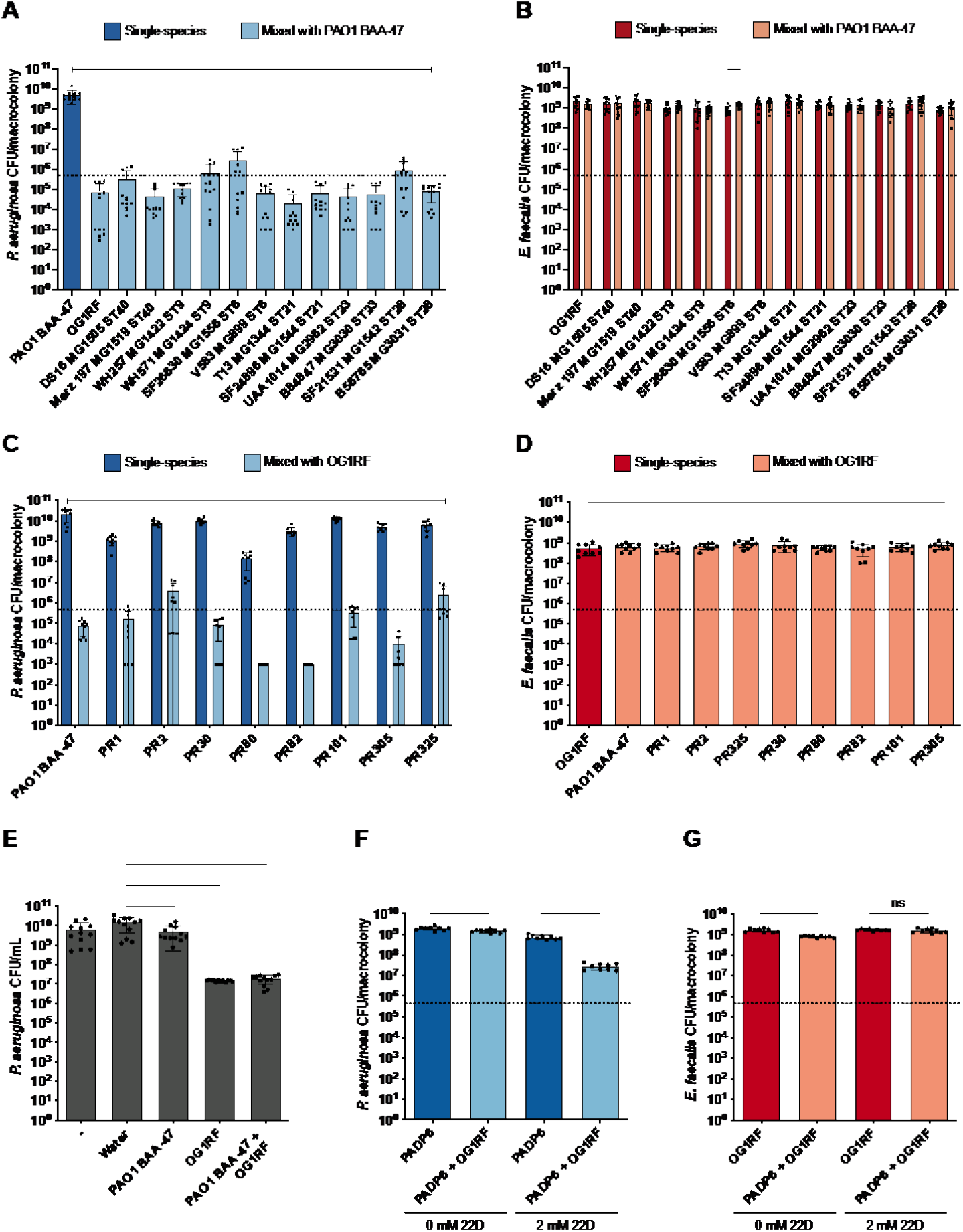
*E. faecalis* inhibits *P. aeruginosa* growth in iron-restricted conditions. Enumeration of **(A)** PAO1 BAA-47, **(B)** *E. faecalis* clinical isolates, **(C)** *P. aeruginosa* clinical isolates and **(D)** OG1RF from 24 h biofilms with single or mixed inoculums grown in media supplemented with 1 mM 22D. Dotted lines represent inoculum of bacteria spotted. N ≥ 3 with 3 technical replicates; error bars represent SD from the mean. Statistical analysis was performed using Mann-Whitney U test, *p < 0.05, **p < 0.01, ***p < 0.001, ****p < 0.0001. Capped line showing statistical significance in **(A)** and **(C)** represents ****p < 0.0001 was observed for all comparisons made. Enumeration of **(E)** PAO1 BAA-47 from 24 h cultures grown with fresh TSBG media, water, or cell-free supernatant obtained from 24 h biofilms of PAO1 BAA-47, OG1RF and PAO1 BAA-47 mixed with OG1RF. The water and respective supernatants were mixed at a 1:1 ratio with fresh TSBG media supplemented with 1 mM 22D for PAO1 BAA-47 growth. N = 4 with 3 technical replicates; error bars represent SD from the mean. Statistical analysis was performed using Mann-Whitney U test, *p < 0.05, **p < 0.01, ***p < 0.001, ****p < 0.0001. Enumeration of **(F)** PADP6 and **(G)** OG1RF from 24 h macrocolonies with single or mixed inoculums grown in media supplemented without and with 2 mM 22D. Bacterial species were mixed at a 1:1 ratio for mixed- species macrocolonies. Dotted lines represent inoculum of bacteria spotted. N = 3 with 3 technical replicates; error bars represent SD from the mean. Statistical analysis was performed using Mann-Whitney U test, *p < 0.05, **p < 0.01, ***p < 0.001, ****p < 0.0001.

We next performed supernatant transfer assays in which either *E. faecalis* or *P. aeruginosa* single- or mixed-species biofilms were grown in iron-restricted media, and their cell-free biofilm supernatants were collected and supplied to *P. aeruginosa* at a 1:1 ratio with fresh media for subsequent growth assays. There were minimal differences in PAO1 BAA-47 growth when supplemented with biofilm supernatant obtained from single-species PAO1 BAA-47 biofilm as compared to supplemented with water (**Figure 1E**). However, when PAO1 BAA-47 was grown with supernatant obtained from single-species OG1RF or mixed PAO1 BAA-47 and OG1RF biofilms, we observed a significant inhibition of PAO1 BAA-46 growth as compared to control supplementation (**Figure 1E**), suggesting that *P. aeruginosa* inhibition is mediated by the presence of *E. faecalis*.

Next, to validate the above findings and to determine the mechanistic basis of polymicrobial interactions between *E. faecalis* and *P. aeruginosa* in iron-restricted conditions, we performed mixed-species macrocolony biofilm assays (57, 58), initially using *E. faecalis* OG1RF and *P. aeruginosa* PAO1 for our experiments. However, PAO1-WT was sensitive to iron restriction at 22D concentrations greater than 1 mM (**Supplementary Figure 1A**) and had a minimum inhibitory concentration (MIC) of 0.8 mM to 22D (data not shown), while there was no effect on OG1RF at this concentration (**Figure 1F**). As such, a *P. aeruginosa* PAO1 spontaneous mutant that was resistant to 22D chelation was generated and named PADP6. Whole genome sequencing of this mutant revealed a single nucleotide polymorphism in the *nalC* repressor gene (4166561G>T, R15L). Although it is unclear how this mutation enhanced 22D resistance in PADP6, mutation in *nalC* causes an overexpression of the iron-regulated *mexAB-oprM* operon encoding the MexAB-OprM efflux pump upon severe iron restriction (59, 60). The mutation in PADP6 restored growth in 2 mM 22D to similar levels as PAO1-WT macrocolonies grown in unchelated media (**Supplementary Figure 1A**). Additionally, we observed no growth defects in either strain in unchelated or at sub-inhibitory concentration of 22D (**Supplementary Figure 1B**). Moving forward, *E. faecalis* OG1RF and *P. aeruginosa* PADP6 were used for all subsequent experiments.

To validate that PADP6 was also susceptible to *E. faecalis*-mediated growth inhibition when iron was restricted (supplemented with 2 mM 22D), we performed the macrocolony biofilm assay and observed that PADP6 growth was inhibited in mixed-species macrocolonies compared to PADP6 single-species macrocolonies (**Figure 1E**), whereas OG1RF growth in the single- and mixed-species macrocolonies was unaffected (**Figure 1F**). Further, the addition of ferric chloride (FeCl3) to 22D-chelated media restored PADP6 growth in the mixed-species macrocolonies to levels similar to PADP6 single-species growth in chelated media without FeCl3 supplementation (**Supplementary Figure 2A and 2B**), indicating that PADP6 growth inhibition in mixed-species macrocolonies is specific to the presence of *E. faecalis* and iron restriction. The supplementation of other trace metals to 22D-chelated media were unable to rescue growth inhibition of *P. aeruginosa* (**Supplementary Figure 2C and 2D**). Hence, we hypothesized that OG1RF produces a factor, or modulates the local environment, such that it is unfavourable for the growth of PADP6 in iron-restricted conditions.

**Figure 2.**
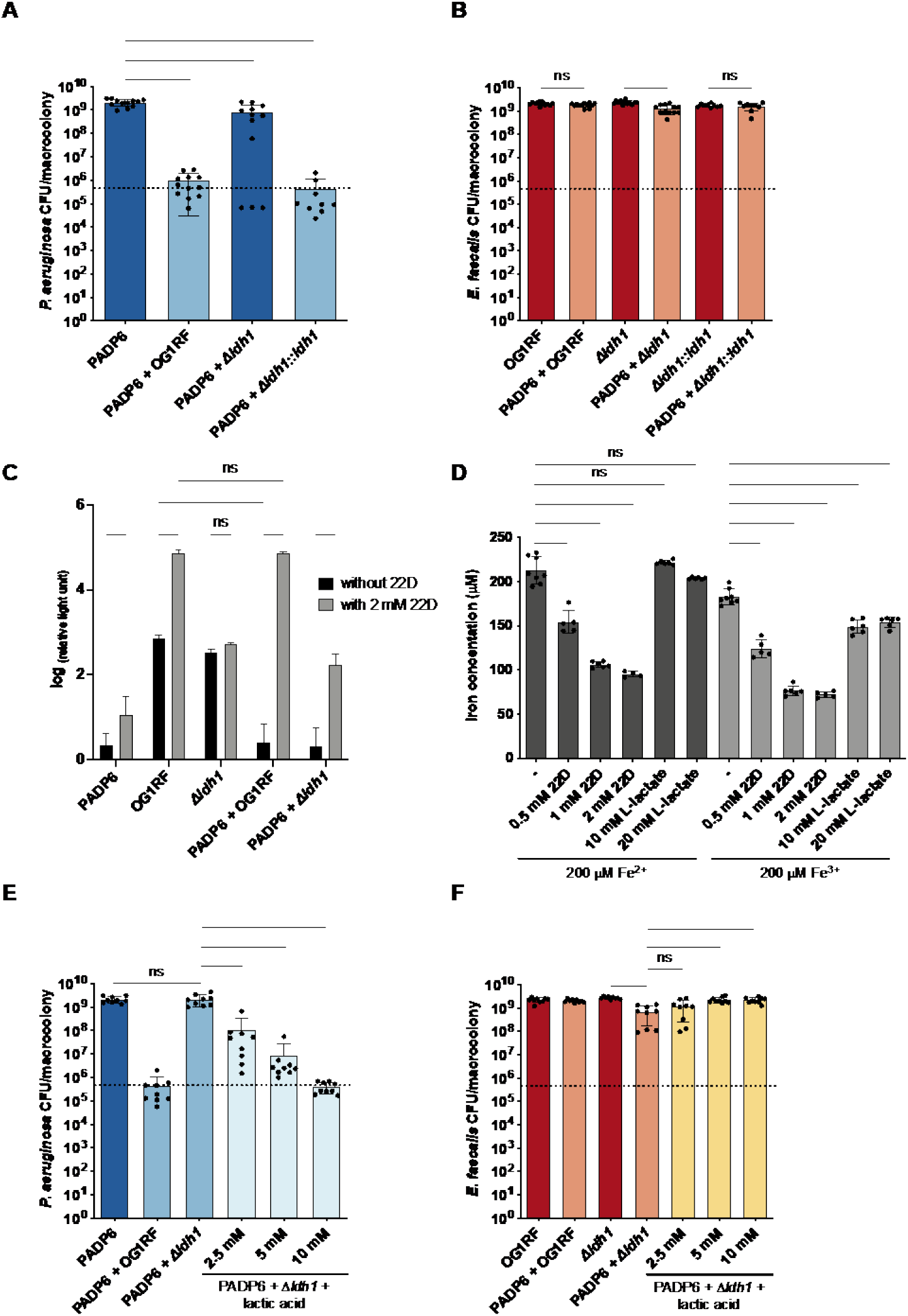
L-lactate produced by *E. faecalis* inhibits *P. aeruginosa* growth in iron- restricted media. Enumeration of **(A)** PADP6 and **(B)** OG1RF, Δ*ldh1*, Δ*ldh1::ldh1* from 48 h macrocolonies with single or mixed inoculums grown in 2 mM 22D-chelated media. Bacterial species were mixed at a 1:1 ratio for mixed-species macrocolonies. Dotted lines represent inoculum of bacteria spotted. N ≥ 3 with 3 technical replicates; error bars represent SD from the mean. Statistical analysis was performed using Mann-Whitney U test, *p < 0.05, **p < 0.01, ***p < 0.001, ****p < 0.0001. **(C)** Quantification of L-lactate exported from 48 h single- and mixed-species macrocolonies grown in media supplemented without and with 2 mM 22D. N = 3 with 2 technical replicates; error bars represent SD from the mean. Statistical analysis was performed using two-way ANOVA with Tukey’s test for multiple comparisons, *p < 0.05, **p < 0.01, ***p < 0.001, ****p < 0.0001. **(D)** Quantification of iron when 200 μM of iron (II) sulfate heptahydrate (Fe^2+^) and iron (III) chloride hexahydrate (Fe^3+^) were supplemented without and with varying concentrations of 22D (0.5, 1 and 2 mM) or L- lactate (10 and 20 mM). N ≥ 3 with 2 technical replicates; error bars represent SD from the mean. Statistical analysis was performed using Mann-Whitney U test, *p < 0.05, **p < 0.01, ***p < 0.001, ****p < 0.0001. Enumeration of **(E)** PADP6 and **(F)** OG1RF, Δ*ldh1* from 48 h macrocolonies with single or mixed inoculums grown in 2 mM 22D- chelated media without and with increasing lactic acid concentrations (2.5, 5, and 10 mM). Bacterial species were mixed at a 1:1 ratio for mixed-species macrocolonies. Dotted lines represent inoculum of bacteria spotted. N = 3 with 3 technical replicates; error bars represent SD from the mean. Statistical analysis was performed using Mann-Whitney U test, *p < 0.05, **p < 0.01, ***p < 0.001, ****p < 0.0001.

Planktonic growth was also examined to determine whether PADP6 growth inhibition in the presence of OG1RF was specific to biofilms. In unchelated media, we observed PADP6 growth inhibition by approximately 1 log in co-culture with *E. faecalis* OG1RF compared to PADP6 alone, while OG1RF growth in co-culture remained unaffected (**Supplementary Figure 3**). However, PADP6 was further inhibited by more than 2 logs in the presence of 22D. Therefore, *E. faecalis* OG1RF inhibition of *P. aeruginosa* PADP6 growth is not a biofilm-specific phenotype.

### *E. faecalis ldh1* is responsible for *P. aeruginosa* growth inhibition in iron- restricted conditions

An *E. faecalis* mariner transposon library screen was performed to identify *E. faecalis* mutants that did not inhibit PADP6 planktonic growth in iron-restricted conditions. We identified six *E. faecalis* mutants that did not inhibit PADP6 growth when co-cultured; however, upon validation of these six mutants in the macrocolony assay, three of which were validated for failure to inhibit PADP6 growth: *ldh1*, *gloA3* and *guaB* (**Supplementary Table 1**). Here we focused on the transposon insertion in *ldh1* (*ldh1::Tn*). In *E. faecalis*, there are two copies of *ldh* (*ldh1* and *ldh2*) that encode L-lactate dehydrogenase (LDH) to catalyse the reduction of pyruvate to L-lactate (61). Of the two copies, *ldh1* accounts for the majority of L-lactate produced in *E. faecalis* in laboratory settings (61, 62).

We then created a *ldh1* deletion mutant (OG1RF Δ*ldh1*) and validated the transposon screening results in a macrocolony biofilm assay with PADP6 and OG1RF Δ*ldh1*. Specifically, PADP6 growth was not inhibited in the mixed PADP6 and OG1RF Δ*ldh1* macrocolonies compared to mixed PADP6 and OG1RF macrocolonies (**Figure 2A**). Importantly, OG1RF Δ*ldh1* growth in single- and mixed-species macrocolonies was unaffected (**Figure 2B**). Upon *ldh1* chromosomal complementation in OG1RF *Δldh1*, PADP6 growth was inhibited to levels as when grown with OG1RF (**Figure 2A**), suggesting that *ldh1* plays a role in inhibiting PADP6 growth in iron-restricted conditions.

### *E. faecalis*-derived L-lactate is responsible for *P. aeruginosa* growth inhibition in mixed-species macrocolonies when iron is restricted

In a previous study, the amount of lactate exported in the supernatant of *E. faecalis* V583 Δ*ldh1* was lower than *E. faecalis* V583 (62). Thus, to investigate the possible role of L-lactate in inhibiting PADP6 growth in mixed-species macrocolonies grown in iron-restriction conditions, we quantified the extracellular L-lactate. We detected significantly more L-lactate from OG1RF single-species macrocolonies grown in iron-restricted compared to unchelated conditions (**Figure 2C**), indicating that OG1RF increased L-lactate production in iron-restricted media is independent of PADP6 presence. Increased L-lactate production was further supported by transcriptomic data, in which we observed upregulation of *ldh1* (log2FC = 0.57) in OG1RF single-species macrocolonies grown in iron-restricted media compared to those grown in unchelated media (**Table 1**). Next, we detected a significant reduction of L-lactate in mixed PADP6 and OG1RF macrocolonies compared to OG1RF single-species macrocolonies in unchelated media, whereas comparable L-lactate levels between single- and mixed-species were detected in iron-restricted conditions (**Figure 2C**). Based on transcriptomic data, we observed an upregulation of *ldh1* (log2FC = 3.27) in mixed PADP6 and OG1RF macrocolonies grown in iron-restricted media compared to those grown in unchelated media (**Table 1**). Together, these data indicate that *ldh1* is upregulated when iron is restricted, leading to increase production and export of L-lactate. In previous studies, lactate was found to chelate iron (63, 64), which we confirmed in our assay conditions for ferric iron (**Figure 2D**). Consequently, even mild iron chelating effects of *E. faecalis*- derived L-lactate could further restrict iron availability in the environment.

**Table 1.** *E. faecalis* L-lactate dehydrogenase differentially regulated in iron-restricted conditions.

| Comparison conditions | Locus tag | Name | Description | log2FC | p-value | FDR |
| --- | --- | --- | --- | --- | --- | --- |
| OG1RF single-species macrocolonies grown in iron restriction relative to OG1RF single-species macrocolonies grown in unchelated conditions | OG1RF_10199 | *ldh1* | L-lactate dehydrogenase | 0.57 | 8.75E-05 | 1.68E-03 |
| Mixed OG1RF and PADP6 macrocolonies grown in iron restriction relative to mixed OG1RF and PADP6 macrocolonies grown in unchelated conditions | OG1RF_10199 | *ldh1* | L-lactate dehydrogenase | 3.27 | 9.02E-25 | 8.91E-24 |

982 *Complete table can be found in the Supplementary file 1.*

To confirm this, we compared PADP6 and OG1RF transcriptomes of mixed PADP6 and OG1RF macrocolonies to mixed PADP6 and OG1RF Δ*ldh1* macrocolonies grown in iron-restricted media. Unfortunately, due to the low PADP6 cell numbers in the mixed PADP6 and OG1RF macrocolonies and therefore, lesser number of PADP6 raw counts, we were unable to draw conclusions regarding iron availability based on the gene expression profile of PADP6 (**Supplementary Table 2**). However, OG1RF iron acquisition genes, such as ABC transporters (65), were upregulated in the mixed PADP6 and OG1RF macrocolonies, suggesting that iron availability was restricted when L-lactate levels was high (**Supplementary Table 3**). Consistent with this, increased L-lactate levels in the environment was inversely correlated with PADP6 growth in the mixed PADP6 and OG1RF macrocolonies in iron-restricted conditions (**Figure 2A and 2C**).

Due to the anionic nature of L-lactate at all metabolic pH, it cannot pass through the *E. faecalis* cell membrane freely (61). As a result, L-lactate is exported out of *E. faecalis* as lactic acid (55) with a pKa of 3.86 (66–68) which is lower than the surrounding environmental pH. Therefore, upon export, lactic acid is deprotonated to L-lactate and releases H^+^ into the environment resulting in a lowered environmental pH. To further investigate the role of L-lactate, we supplemented increasing amounts of lactic acid to mixed PADP6 and OG1RF Δ*ldh1* macrocolonies in 22D-chelated media. We observed a dose-dependent inhibition of PADP6 growth in the mixed-species macrocolonies with increasing lactic acid concentrations from 2.5 mM to 10 mM (**Figure 2E**), while OG1RF Δ*ldh1* growth in the mixed-species macrocolonies remained relatively unchanged (**Figure 2F**).

### L-lactate is necessary, but not sufficient, for inhibiting *P. aeruginosa* growth in iron-restricted conditions

To investigate whether L-lactate alone was sufficient for inhibiting PADP6 growth in iron-restricted media, we grew PADP6 single-species macrocolonies supplemented with increasing 22D and lactic acid concentrations. When supplemented with 10 mM or 20 mM lactic acid in unchelated media, PADP6 growth remained relatively unchanged compared to unchelated media without lactic acid supplementation (**Figure 3A**). Based on the lactic acid supplementation results obtained in **Figure 2E and 2F**, we expected that PADP6 growth would be inhibited when supplemented with 10 mM lactic acid in iron-restricted conditions. However, upon supplementation of 10 mM lactic acid, we only observed a significant inhibition of PADP6 growth when it was grown in 4 mM 22D-chelated media, but not in 2 mM and 3 mM 22D-chelated media as compared to unchelated media (**Figure 3A**). Whereas, when supplemented with 20 mM lactic acid, we observed significant PADP6 growth inhibition at all tested 22D concentrations, as compared to unchelated media (**Figure 3A**). A possible explanation for why more lactic acid was needed to inhibit single-species PADP6 growth in 2 mM 22D could be that *E. faecalis ldh2* is also contributing to L-lactate production in OG1RF Δ*ldh1*, hence a lower amount of lactic acid was sufficient to inhibit PADP6 growth in the mixed-species macrocolonies.

**Figure 3.**
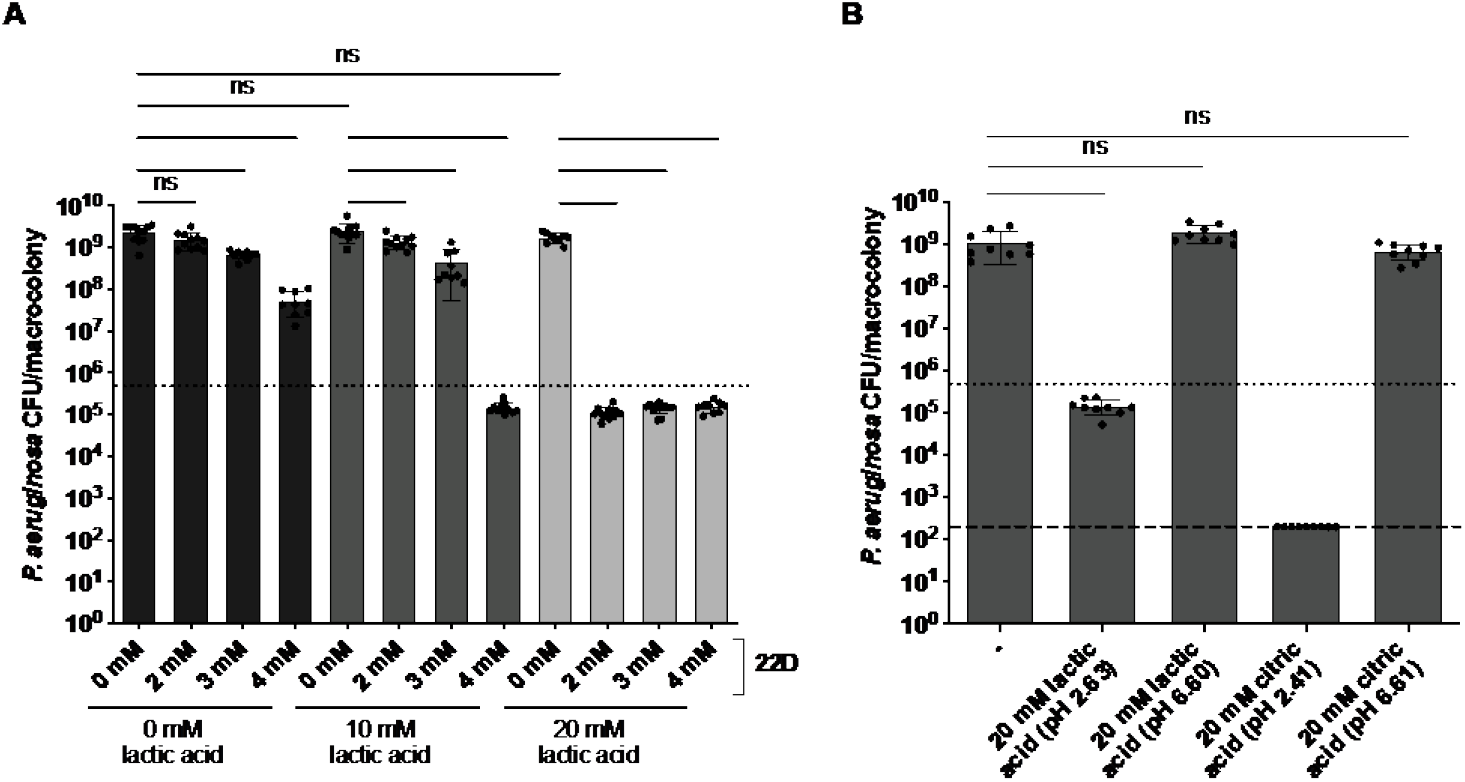
*P. aeruginosa* growth inhibition is due to lowered environmental pH in iron-restricted conditions. **(A)** Enumeration of PADP6 from 48 h single-species macrocolonies grown in media supplemented without and with increasing 22D concentrations (2, 3 and 4 mM), which is then further supplemented without and with lactic acid (10 and 20 mM). Dotted lines represent inoculum of bacteria spotted. N ≥ 3 with 3 technical replicates; error bars represent SD from the mean. Statistical analysis was performed using Mann-Whitney U test, *p < 0.05, **p < 0.01, ***p < 0.001, ****p < 0.0001. **(B)** Enumeration of PADP6 from 48 h single-species macrocolonies grown in 2 mM 22D-chelated media supplemented without and with 20 mM lactic acid (pH unadjusted and pH adjusted to pH 6.6) or 20 mM citric acid (pH unadjusted and pH adjusted to pH 6.61). Dotted lines represent inoculum of bacteria spotted and dashed lines represent limit of detection. N = 3 with 3 technical replicates; error bars represent SD from the mean. Statistical analysis was performed using Mann-Whitney U test, *p < 0.05, **p < 0.01, ***p < 0.001, ****p < 0.0001.

Taken together, these data demonstrate that L-lactate is necessary, but not sufficient, for PADP6 growth inhibition in iron-restricted media.

### Decreased environmental pH under iron restriction inhibits *P. aeruginosa* growth

Next, we investigated why PADP6 viability was lost in the presence of L-lactate in iron-restricted conditions. Upon export of lactic acid by *E. faecalis*, it is deprotonated into L-lactate and H^+^. We therefore examined whether a lowered environmental pH contributes to PADP6 growth inhibition when iron was restricted. Supplementation of iron-restricted media with 20 mM pH unadjusted lactic acid (pH 2.69) resulted in significant PADP6 growth inhibition as compared to the absence of lactic acid supplementation (**Figure 3B**). In contrast, PADP6 growth was unaffected when we supplemented the media with 20 mM pH adjusted lactic acid (pH 6.60) using sodium hydroxide (NaOH) when iron was otherwise restricted (**Figure 3B**). To examine if other organic acids had the ability to inhibit *P. aeruginosa* growth, we also supplemented the iron-restricted media with citric acid to lower the environmental pH. Supplementation with pH unadjusted citric acid (pH 2.41) significantly inhibited PADP6 growth to below the limit of detection, while no significant growth difference was observed upon supplementation with 20 mM pH adjusted citric acid using NaOH (pH 6.61) or in the absence of citric acid supplementation (**Figure 3B**). Moreover, alleviation of the low pH in iron-restricted media with PIPES, MOPS or HEPES buffer partially rescued *P. aeruginosa* growth in the mixed-species macrocolonies (**Supplementary figure 4**). Altogether, these data show that the lowered environmental pH as a consequence of *E. faecalis* lactic acid export, coupled with L-lactate-mediated chelation of iron, plays a critical role in *P. aeruginosa* growth in iron-restricted conditions.

**Figure 4.**
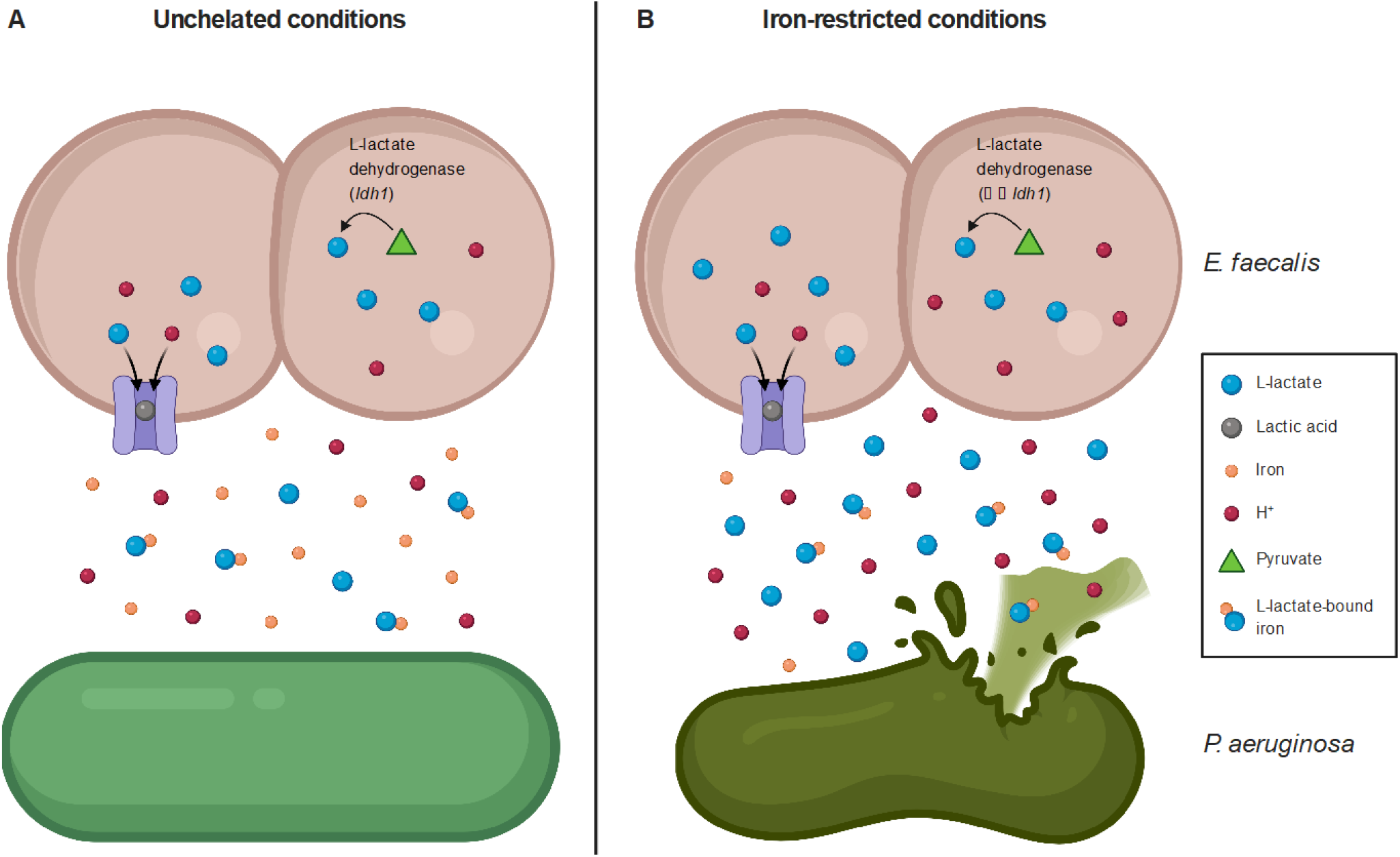
Proposed working model of *E. faecalis* and *P. aeruginosa in vitro* polymicrobial interactions. Interactions between *E. faecalis* and *P. aeruginosa* in **(A)** unchelated and **(B)** iron-restricted conditions. **(A)** In unchelated conditions, L-lactate produced in *E. faecalis* is exported with hydrogen ions via a symporter (purple) as lactic acid, which is then deprotonated in the environment into L-lactate and hydrogen ions (H^+^). These L-lactate then chelates iron in the environment. **(B)** In iron-restricted conditions, *E. faecalis ldh1* expression is upregulated. Consequently, as *E. faecalis* grows, L-lactate production and lactic acid secretion increases. This further chelates iron in an iron-restricted conditions and lowers the environmental pH to a point that *P. aeruginosa* could not grow in.

## DISCUSSION

In this study, we sought to characterize polymicrobial interactions between commonly co-isolated *E. faecalis* and *P. aeruginosa* in an infection-relevant, iron-restricted condition. We show that *E. faecalis* inhibits *P. aeruginosa* growth when iron availability is restricted in an LDH-dependent manner. Since *E. faecalis* LDH1 catalyses the reduction of pyruvate to L-lactate, the increased *ldh1* expression translates to an increased L-lactate production in *E. faecalis*. The L-lactate produced is then exported as lactic acid which then gets deprotonated into L-lactate releasing H^+^ in the surrounding environment. Together, the chelation of iron by L-lactate and lowered pH environment contributes to *P. aeruginosa* growth inhibition in iron-restricted conditions.

Although we showed that *E. faecalis* antagonize *P. aeruginosa* in iron-restricted conditions *in vitro*, they are often co-isolated *in vivo* (48, 50, 51, 53). During *P. aeruginosa* wound infections, genes involved in siderophore biosynthesis are upregulated, suggesting that iron availability is indeed restricted during wound infection (69). Therefore, our *in vitro* assay model is infection-relevant and the contrasting observation that we observed *in vitro* and *in vivo* could be due to in-host bacterial or host mechanism that negate or override the antagonism. Our findings show that, in addition to iron-restrictive effects, *P. aeruginosa* growth inhibition is largely dependent on the effects of pH arising from export of *E. faecalis* lactic acid into the environment. As such, co-existence of *E. faecalis* and *P. aeruginosa in vivo* may be due to host influence on *E. faecalis* L-lactate production, or that the host buffers the environmental pH negating *E. faecalis* L-lactate effects (70). Another possibility could be differences in *in vitro* and *in vivo* spatial organization of *E. faecalis* and *P. aeruginosa*. When *P. aeruginosa* and *E. faecalis* were grown *in vitro*, they exhibit distinct spatial separation in which *P. aeruginosa* forms a structured biofilm above the *E. faecalis* biofilm (71). Whereas during polymicrobial wound infection (*P. aeruginosa*, *E. faecalis*, *S. aureus*, and *F. magna*), *P. aeruginosa* is seen throughout the wound bed as well as at the leading edge of wound (52). Host factors might also contribute and affect spatial organization as this is evident in gut microbiome spatial organization (72, 73). Spatial structuring that keeps the two species physically separated could blunt any local pH and iron-competition effects.

An outstanding question from this work is how iron restriction leads to an upregulation of *E. faecalis ldh1*. *E. faecalis* possesses two copies of the *ldh* gene, *ldh1* and *ldh2*, both encoding for L-lactate dehydrogenase (61). Both isoenzymes contributes to L- lactate production through catalysing the reduction of pyruvate to L-lactate (61). The activity of both isoenzymes is regulated by fructose 1,6-bisphosphate, intracellular phosphate and pH levels (74). The principal L-lactate dehydrogenase, encoded by *ldh1*, was suggested to be post-transcriptionally regulated upon different growth rates (75). The transcription of *ldh1* is also activated by CcpA, a global transcription regulator of carbon catabolite repression by binding to a catabolite-responsive elements (*cre*) box identified upstream of *ldh1* gene (76). However, little is known if iron levels play any role in regulating *ldh1* expression. We show that *E. faecalis ldh1* expression is increased in iron-restricted media and this is supported by an increased amount of L-lactate measured in the surrounding environment of macrocolonies grown. Ferric uptake regulator (Fur) is a transcription factor involved in regulating iron uptake and homeostasis (77). The DNA-binding sequence of Fur is well-studied (65, 78, 79) and we tried using different DNA Fur binding motifs to identify possible Fur binding sites upstream of *ldh1*, but we did not find any that resemble known binding motifs. Despite this, a previous study reported that *ldh1* is differentially expressed between *E. faecalis* OG1RF and OG1RF Δ*fur* mutant, suggesting that *ldh1* expression is directly or indirectly influenced by Fur (65). An upregulation of *ldh1* and several genes involved in iron transport is also observed in another transcriptome study done when *E. faecalis* was exposed to urine (80). It is therefore consistent that in iron-restricted environments, such as urine or *E. faecalis* macrocolonies growing in iron-restricted media, *ldh1* expression is induced. This observation is not limited to *E. faecalis* as *ldh1* expression is similarly induced in iron-restricted conditions for the anaerobe *Clostridium acetobutylicum* (81). Although it remains unclear how iron levels influence *ldh1* expression in *E. faecalis*, it is tempting to speculate that there is an interplay between iron levels and energy metabolism during growth in iron-restricted conditions.

Lactic acid exported by *E. faecalis* is deprotonated to L-lactate and H^+^. We show that the consequential lowered environmental pH contributes to *P. aeruginosa* growth inhibition when iron is restricted. This is not surprising as *P. aeruginosa* growth is generally affected at low pH and *P. aeruginosa* prefers to grow in a more neutral pH range (82, 83). In fact, *E. faecalis* V583 adopts a similar strategy of lowering environmental pH as a result of lactic acid export to inhibit *Klebsiella pneumoniae* growth in polymicrobial biofilms (84). Even though a *P. aeruginosa* PADP6 strain that is able to grow in iron-restricted conditions is used to study the polymicrobial interactions with *E. faecalis*, the lowered iron availability resulted from 22D-mediated iron chelation and L-lactate-mediated iron chelation is likely an added stress apart from the lowered pH environment. Afterall, iron is an essential element for many cellular and metabolite processes (85, 86). Moreover, the low pH might also affect the binding affinity of iron to *P. aeruginosa* siderophores. The binding affinity for iron and zinc of *E. coli* Nissle siderophore yersiniabactin changes accordingly to pH, in which yersiniabactin preferentially binds to zinc as pH increases (87). Hence, the low pH environment created by *E. faecalis* might possibly affect *P. aeruginosa* siderophore-mediated iron uptake and consequently, negatively impacting its growth due to insufficient intracellular iron.

It remains unclear how *nalC* mutation in PADP6 which overexpresses MexAB-OprM efflux pump confers tolerance to iron starvation (59). Although *P. aeruginosa* PvdRT-OpmQ (88, 89) and MexAB-OprM (60) efflux pumps are implicated in pyoverdine secretion, recent research suggests that MexAB-OprM efflux pump is not involved (89). An alternative hypothesis is that PADP6 might have a compromised outer membrane. In *E. coli*, Δ*tonB* mutants were found to grow poorly in iron-restricted media and prolonged incubation resulted in mutations in *yejM* gene which encodes for a putative inner to outer membrane cardiolipin transporter (90). The transport of iron chelator, enterobactin, back into the cell is through outer membrane receptor protein FepA, which activity is dependent on energy transduced from proton motive force by TonB- ExbDB complex (91). Further characterization revealed that Δ*tonB* mutants with mutations in *yejM* are hypersensitive to vancomycin, suggesting that its outer membrane permeability is compromised to overcome the iron-restricted environment (90). These observations suggest that by having mutations in *yejM*, at the cost of having a more permeable membrane, could allow transport of enterobactin into the cell without FepA. Similarly, PADP6 might have a compromised outer membrane which facilitates the transport of pyoverdine and pyochelin into itself.

Based on the current findings, we propose a working model of *E. faecalis* and *P. aeruginosa* polymicrobial interactions *in vitro* (**Figure 4**). During co-culture of *E. faecalis* and *P. aeruginosa* in unchelated media, L-lactate produced by *E. faecalis* is exported out via a symporter with H^+^ as lactic acid. Since the pKa of lactic acid is lower than pH of the environment, it gets deprotonated into L-lactate and H^+^, acidifying the environment. The L-lactate in the environment then chelates iron in the media (**Figure 4A**). In contrast, during co-culture in iron-restricted conditions, there is an upregulation of *E. faecalis ldh1* expression which translates to an increased L-lactate production and lactic acid exported, further acidifying the environment. As *E. faecalis* continues to grow and lactic acid accumulates over time, the L-lactate in the environment further restricts iron availability in the iron-restricted media and the acidity of the environment subsequently exceeds a pH threshold at which *P. aeruginosa* can no longer grow (**Figure 4B**).

Many infections are often polymicrobial and our work highlights the importance of how changes in microenvironment such as iron, pH levels, or host-related factors can significantly influence the interactions between two bacterial species. Knowledge of such *in vitro* antagonism between species could potentially be used as a basis for additional control strategies against specific bacterial pathogens in the management of infections.

## MATERIALS AND METHODS

### Bacterial strains and growth conditions

Bacterial strains used in this study are listed in **Supplementary Table 4**. Unless stated, all *P. aeruginosa* and *E. faecalis* bacterial strains were grown at 37 °C in shaking or static conditions for 16 – 18 h, respectively. Cells were harvested by centrifugation at 12,000 *g* for 5 min and cell pellets were washed twice with 1 mL of 1X sterile phosphate buffered saline (PBS). The final pellet was resuspended in 3 mL of 1X sterile PBS prior to optical density (OD) measurement at 600 nm. Cell suspensions were then normalized to the required cell number for different experimental assays. For *Pseudomonas* selection, bacteria were spotted onto *Pseudomonas* Isolation agar (PIA) (Difco™ BD, USA) supplemented with 100 µg/mL ampicillin (Sigma-Aldrich, USA). For *E. faecalis* OG1RF selection, bacteria were spotted onto Tryptone Soya Broth (TSB) (Oxoid, Canada) solidified with 1.5% agar (Oxiod Technical No. 3) and supplemented with 10 mM glucose (TSBG), 10 µg/mL colistin (Sigma-Aldrich, USA) and 10 µg/mL nalidixic acid (Sigma-Aldrich, USA), respectively.

### Planktonic, static biofilm and macrocolony biofilm assays

Bacterial cultures were normalized to 1 – 2 × 10^8^ CFU/mL in 1X PBS. Macrocolonies were produced by inoculating 5 μL of the respective bacterial cultures onto the surface of TSBG solidified with 1.5% agar and incubated at 37 °C for either 24 h or 48 h. For mixed-species macrocolonies, bacterial species were mixed at a 1:1 ratio. When appropriate, the TSBG agar was further supplemented with or without 2,2’ Bipyridyl (22D) (Sigma-Aldrich, USA), iron (III) chloride hexahydrate (Merck, USA), citric acid (Merck, USA), lactic acid, iron (II) sulfate heptahydrate, copper (II) chloride anhydrous, manganese (II) sulfate, magnesium (II) chloride, zinc chloride anhydrous, 1,4- Piperazinediethanesulfonic acid disodium salt (PIPES), 4-Morpholinepropanesulfonic acid, 3-(N-Morpholino)propanesulfonic acid (MOPS) or 4-(2-hydroxyethyl)-1- piperazineethanesulfonic acid (HEPES) buffer (all purchased from Sigma-Aldrich, USA). Macrocolonies were excised and resuspended in 2 mL of sterile 1X PBS, followed by bacteria enumeration on respective selection agar. For supernatant transfer and static biofilm assays, single- and mixed-species inocula were prepared as described above and inoculated in TSBG media for 24 h at 37 °C unless stated otherwise. For static biofilm assays, CFU enumeration were performed first by scraping the wells of 6-well microtiter plates, then pipetting to mix homogenously and serially diluted for plating on selective agar plates. For preparation of supernatant media, 24 h biofilms were first scraped, suspended in conical tubes (cells and spent media together) and centrifuged at 4,000 rpm for 20 min to pellet the cells. Spent media is transferred to a new tube and filter-sterilized to obtain cell-free supernatant. The cell-free supernatant was then mixed with fresh TSBG media or water, and then supplemented with 1 mM 22D, prior to inoculating for subsequent growth at 37 °C in static conditions for 24 h. For planktonic assay, 5 μL of the respective bacterial cultures were inoculated into TSBG media and incubated at 37 °C in shaking conditions for 24 h.

### Construction of PADP6 and PADP6-mCherry strains

Overnight cultures of PAO1- WT were diluted to 10^9^, 10^8^, 10^7^ and 10^6^ CFU/mL in 1X PBS, and 300 µL of each cell suspension were plated onto LB Lennox agar (Difco™ BD, USA) supplemented with 1.5, 2, 2.5, 3 and 4 mM 22D. Plates were incubated at 37 °C for 24 to 36 h. PAO1-WT and few colonies that grew in 3 mM 22D-chelated condition were isolated and gDNA extracted using Wizard® Genomic DNA Purification Kit (Promega, USA) for use in whole genome sequencing. PADP6 was chromosomally tagged with mCherry through triparental conjugation using PADP6 as a recipient with delivery plasmid pUC18- miniTn7-P*tac*-mCherry (*E. coli*) and helper plasmid pTNS1 (*E. coli*) resulting in PADP6- mCherry (92–94).

### Genome sequencing and analysis

Raw reads were imported into CLC Genomics Workbench 8.0 (Qiagen, Germany), followed by quality trimming to remove bad quality reads. The trimmed reads were then mapped to the reference genome before the Basic Variant Detection module was used to detect for mutations using the default parameters. The mutations detected in the isolate was then filtered against PAO1-WT control to determine the mutations acquired for survival in iron-restricted conditions.

### Planktonic growth assay

Bacterial cultures were normalized to OD600 of 0.01 in the respective media and inoculated into 24-well microtiter plates. All microtiter plates were incubated at 37 °C in shaking conditions. Planktonic growth was measured by recording the OD600 between 30 min to 1 h intervals using a Tecan Infinite© M200 Pro spectrophotometer (Tecan Group Ltd., Switzerland) until early stationary growth phase was reached.

### *E. faecalis* transposon library screen

An *E. faecalis* OG1RF mariner transposon library consisting of 14,978 mutants was cryogenically stocked in 96-well microtiter plates (95). These OG1RF transposon mutants were cultured in 180 µL BHI broth at 37 °C in static conditions for 16 – 18 h in 96-well microtiter plates using a cryo-replicator (Adolf Kühner AG, Switzerland) and spotted onto BHI agar plates for incubation at 37 °C for 24 h. Following that, OG1RF transposon mutants from the BHI agar plates were cultured for primary screening in 180 µL BHI broth as described above. PADP6-mCherry cultures were grown and washed as described above. Both the OG1RF transposon mutant cultures and PADP6-mCherry were normalized to OD600 of 0.01 in TSBG media supplemented with 1.2 mM 22D. A primary screen of the *E. faecalis* transposon library was done by mixing the normalized OG1RF transposon mutant cultures and PADP6-mCherry at a 1:1 ratio in 96-well microtiter plates (total volume of 200 µL). The microtiter plates were then incubated at 37 °C in static conditions for 22 h. The growth of PADP6-mCherry was quantified by measuring mCherry fluorescence intensity (excitation = 480 nm, emission = 615 nm) using a Tecan Infinite© M200 Pro spectrophotometer. Secondary validation of the OG1RF transposon mutants was performed by mixed-species macrocolony biofilm assays as described above to quantify the growth of PADP6 and each transposon mutant (CFU/mL) in 1 mM 22D iron-restricted conditions at 37 °C for 24 h.

### Molecular cloning

The primers used in this study are listed in **Supplementary Table 5**. Transformants were screened using respective selection agar as follows: (A) *E. coli* strains, LB with 500 µg/mL erythromycin (pGCP213); and (B) *E. faecalis* strains, BHI with 25 µg/mL erythromycin (pGCP213). Generation of *E. faecalis* knock-out mutants were done by allelic replacement using a temperature-sensitive shuttle vector described previously (96). Vector pGCP213 was linearized using restriction enzymes (New England Biolabs, USA) for the construction of OG1RF *Δldh1* and OG1RF *Δldh1::ldh1*. Linearized vector and inserts were ligated using In-Fusion® HD Cloning Kit (Clontech, Takara, Japan) and transformed into Stellar^TM^ competent cells. Successful plasmid constructs were verified by Sanger sequencing and subsequently extracted and transformed into OG1RF. Transformants were selected with erythromycin at 30 °C, then passaged at non-permissive temperature at 42 °C with erythromycin to select for bacteria with successful plasmid integration into the chromosome. For plasmid excision, bacteria was serially passaged at 37 °C without erythromycin for erythromycin-sensitive colonies. These colonies were then subjected to PCR screening for detection of deletion mutant (OG1RF *Δ*OG1RF_10021) or chromosomal complementation of *ldh1* (OG1RF *Δldh1::ldh1*).

### Lactate-Glo™ assay

L-lactate quantification was done using the Lactate-Glo™ assay kit (Promega, USA). Macrocolonies were excised, resuspended in 5 mL of sterile 1X PBS and homogenized. Supernatant were then collected by centrifuging homogenate at 5,000 *g* for 10 min and used for L-lactate quantification. Briefly, an equal volume of Lactate Detection Reagent was added to the supernatant, and incubated for 60 min at room temperature before luminescence was read using a Tecan Infinite© M200 Pro spectrophotometer.

### Total iron quantification

Iron quantification was done using the Iron Assay Kit (Colorimetric) (Abcam, UK) as per manufacturer’s instruction. Prior to quantification, samples were prepared by supplementing different concentrations of 22D or sodium L-lactate (Sigma-Aldrich, USA) to 200 μM iron (II) sulfate heptahydrate (FeSO4·7H2O) and iron (III) chloride hexahydrate (FeCl3·6H2O). The output was measured immediately at OD593 using a Tecan Infinite© M200 Pro spectrophotometer. The iron concentration in each sample was computed based on the standard curve generated using the iron standards.

### RNA extraction from macrocolonies

Matured single and mixed-species macrocolonies grown for 48 h on TSBG agar supplemented with and without 2,2’ bipyridyl, in biological triplicates, were first scraped into RNAprotect™ Bacteria Reagent (Qiagen, Germany) and incubated at room temperature for 5 mins before centrifuging at 10,000 *g* for 10 mins. The supernatant was decanted and bacteria pellets collected were then subjected to total RNA extraction using RNeasy® mini kit (Qiagen, Germany) with slight modifications. Briefly, cell pellets were resuspended in TE buffer containing 20 mg/mL lysozyme (Sigma-Aldrich, USA) and each sample was further supplemented with 20 µL proteinase K (Qiagen, Germany). This was followed by incubation at 37 °C for 1 h, and subsequent extraction steps was performed according to manufacturer’s protocol. Extracted RNA samples were treated with DNase (TURBO DNA-free™ kit, Invitrogen, USA) for removal of contaminating genomic DNA before it was purified with Monarch® RNA cleanup kit (New England Biolabs, USA). The concentration of RNA and potential DNA contamination were quantified using Qubit™ RNA BR and Qubit™ dsDNA HS assay kits, respectively (Invitrogen, USA). The extracted RNA was quality checked using a TapeStation instrument (RNA ScreenTape, Aligent Technologies, USA) before it was sent for sequencing. Every sample had to have a minimum RNA concentration of 40 – 80 ng/µL, a maximum of 10% DNA contamination and a RINe value ≥ 8.0, before being used for library preparation and subsequent sequencing as 100 bp paired end reads on an Illumina HiSeq2500 at Singapore Centre for Environmental Life Sciences Engineering (SCELSE) sequencing facility.

### Transcriptomic analysis

The raw reads obtained were checked using FastQC (Version 0.11.9) and adaptor trimmed using bbduk from BBMap tools (Version 39.79) (97). Trimmed reads were then mapped using bwa-mem of BWA (Version 0.7.17- r1188) with options “-T 20 -k 13” against *E. faecalis* OG1RF (NCBI accession: CP002621) or *P. aeruginosa* PAO1 (NCBI accession: NC_002516) reference genomes. Reads mapped to predicted open reading frames were quantified using htseq-count of HTSeq (Version 0.12.4) with option “-m intersection-strict” (98). Ribosomal sequences were filtered out from all data sets. Differential gene expression analysis was performed in R using *edgeR* (Version 3.28.1) (99). The log2 fold change values extracted were based on the false discovery rate (FDR) < 0.05.

### Statistical analysis

Statistical analyses were performed with GraphPad Prism software (Version 9.0.0, California, USA) and are described in the respective figure legends.

### Data availability

All RNA-seq sequences were deposited in the National Center for Biotechnology Information Gene Expression Omnibus database under accession number GSE190090.

## ACKNOWLEDGEMENTS

This work was supported by the National Research Foundation and Ministry of Education Singapore under its Research Centre of Excellence Programme, by the Singapore Ministry of Education under its Tier 2 program (MOE2014-T2-1-129) awarded to K.A.K., and by NIAID R21 AI37446 to J.A.L. Preparation of this article was also financially supported by the Interdisciplinary Graduate Programme of Nanyang Technological University. We thank Yang Liang from Southern University of Science and Technology for the *P. aeruginosa* PAO1-WT and *E. coli* strains, Sam P. Brown from Georgia Institute of Technology for the *P. aeruginosa* clinical isolates and Michael S. Gilmore from Harvard Medical school for the *E. faecalis* clinical isolates. Figure 4 in this article was created with BioRender.com.

## CONFLICT OF INTEREST

The authors declare that they have no conflicts of interest with the contents of this article.

---

**Supplementary Figure 1.**
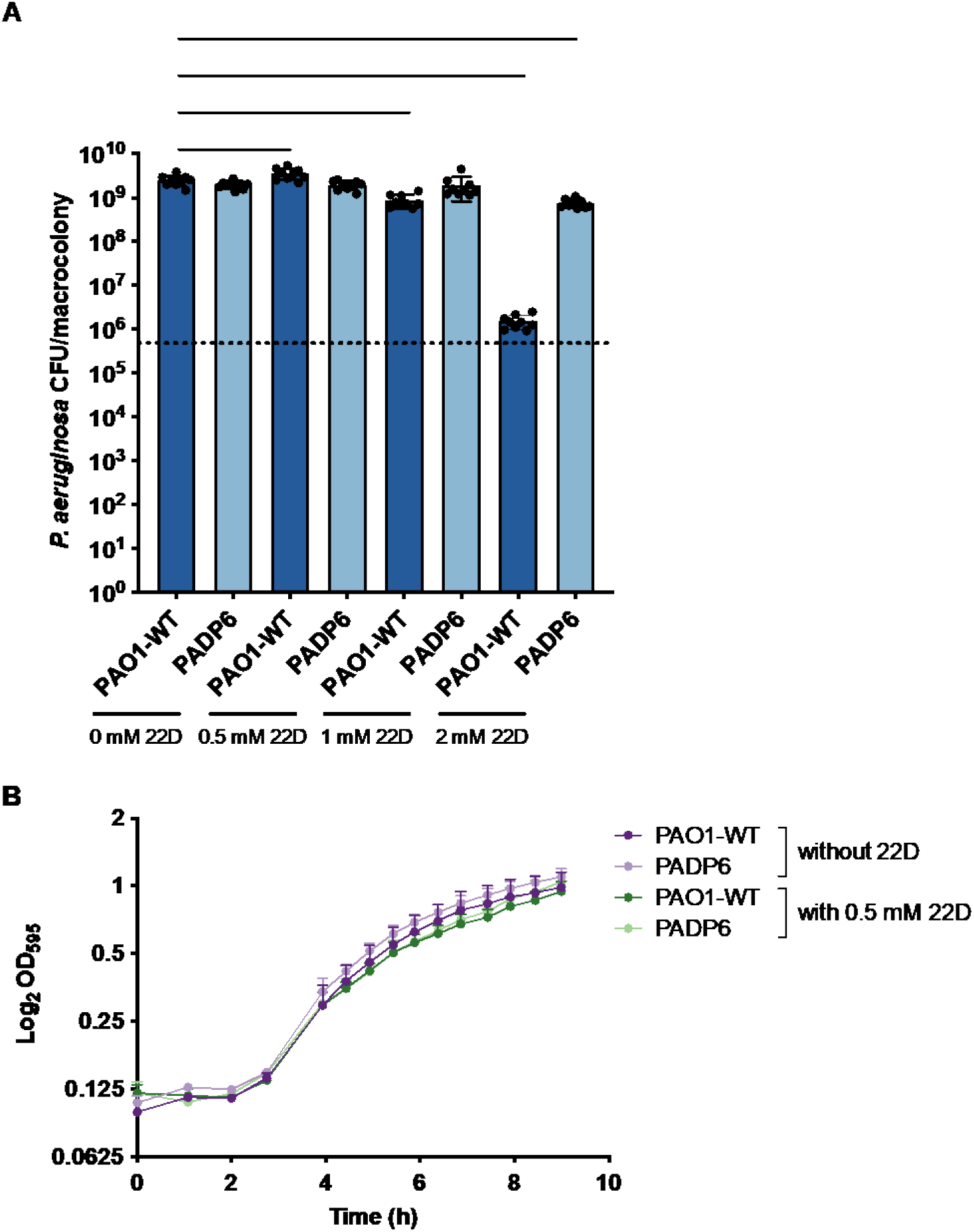
*P. aeruginosa* growth responses to iron restriction. **(A)** Enumeration of PAO1-WT and PADP6 from 24 h single-species macrocolonies grown in media supplemented without and with increasing 22D concentrations (0.5, 1 and 2 mM). Dotted lines represent inoculum of bacteria spotted. N = 3 with 3 technical replicates; error bars represent SD from the mean. Statistical analysis was performed using Mann-Whitney U test, *p < 0.05, **p < 0.01, ***p < 0.001, ****p < 0.0001. **(B)** Planktonic growth of PAO1-WT and PADP6 in TSBG media supplemented without and with 0.5 mM 22D. N = 3 with 3 technical replicates; error bars represent SD from the mean.

**Supplementary Figure 2.**
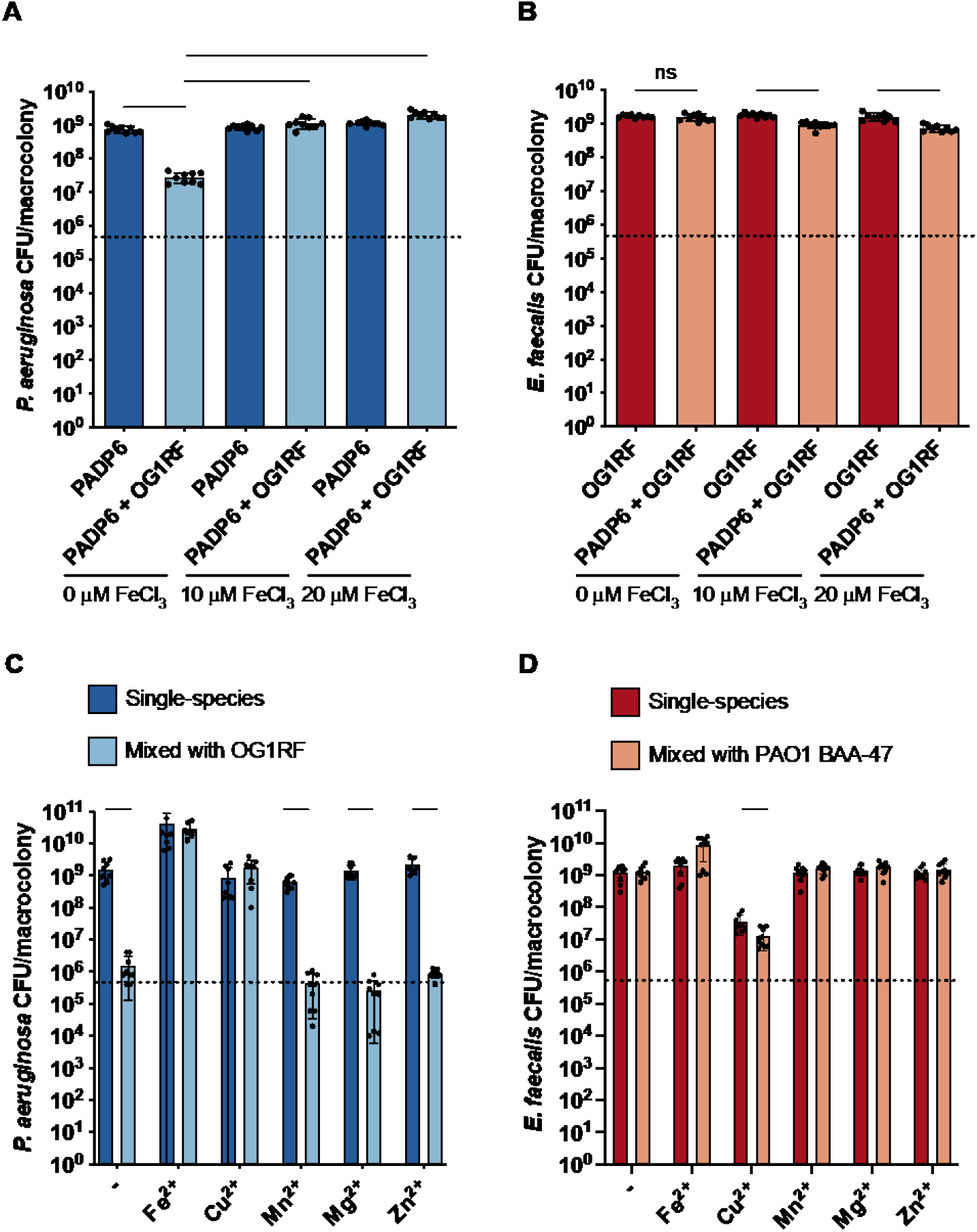
Growth inhibition of *P. aeruginosa* by *E. faecalis* is iron-specific. Enumeration of **(A)** PADP6 and **(B)** OG1RF from 24 h macrocolonies with single or mixed inoculums grown in 2 mM 22D-chelated media without and with FeCl3 (10 and 20 μM). Bacterial species were mixed at a 1:1 ratio for mixed-species macrocolonies. Dotted lines represent inoculum of bacteria spotted. N = 3 with 3 technical replicates; error bars represent SD from the mean. Statistical analysis was performed using Mann-Whitney U test, *p < 0.05, **p < 0.01, ***p < 0.001, ****p < 0.0001. Enumeration of **(C)** PAO1 BAA-47 and **(D)** OG1RF from 24 h single- and mixed-species macrocolonies grown in 1 mM 22D-chelated media without and with 100 μM of varying trace metals. Bacterial species were mixed at a 1:1 ratio for mixed-species macrocolonies. Dotted lines represent inoculum of bacteria spotted. N = 3 with 3 technical replicates; error bars represent SD from the mean. Statistical analysis was performed using Mann-Whitney U test, *p < 0.05, **p < 0.01, ***p < 0.001, ****p < 0.0001.

**Supplementary Figure 3.**
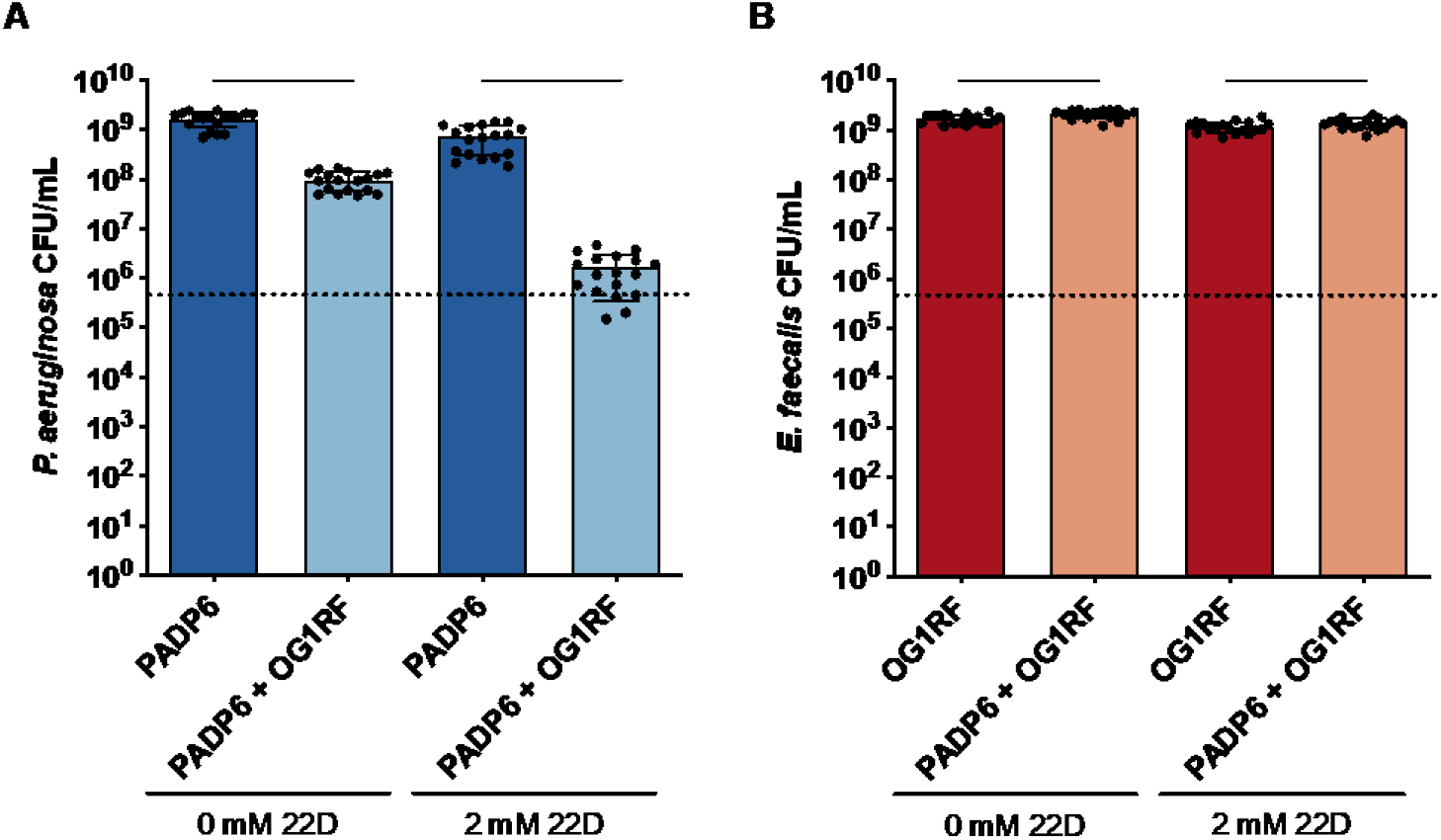
*E. faecalis* inhibits *P. aeruginosa* growth in planktonic conditions regardless of iron levels. Enumeration of **(A)** PADP6 and **(B)** OG1RF from single or mixed inoculums grown for 24 h in media supplemented without and with 2 mM 22D. Bacterial species were mixed at a 1:1 ratio for mixed-species macrocolonies. Dotted lines represent inoculum of bacteria spotted. N ≥ 3 with 3 technical replicates; error bars represent SD from the mean. Statistical analysis was performed using Mann-Whitney U test, *p < 0.05, **p < 0.01, ***p < 0.001, ****p < 0.0001.

**Supplementary Figure 4.**
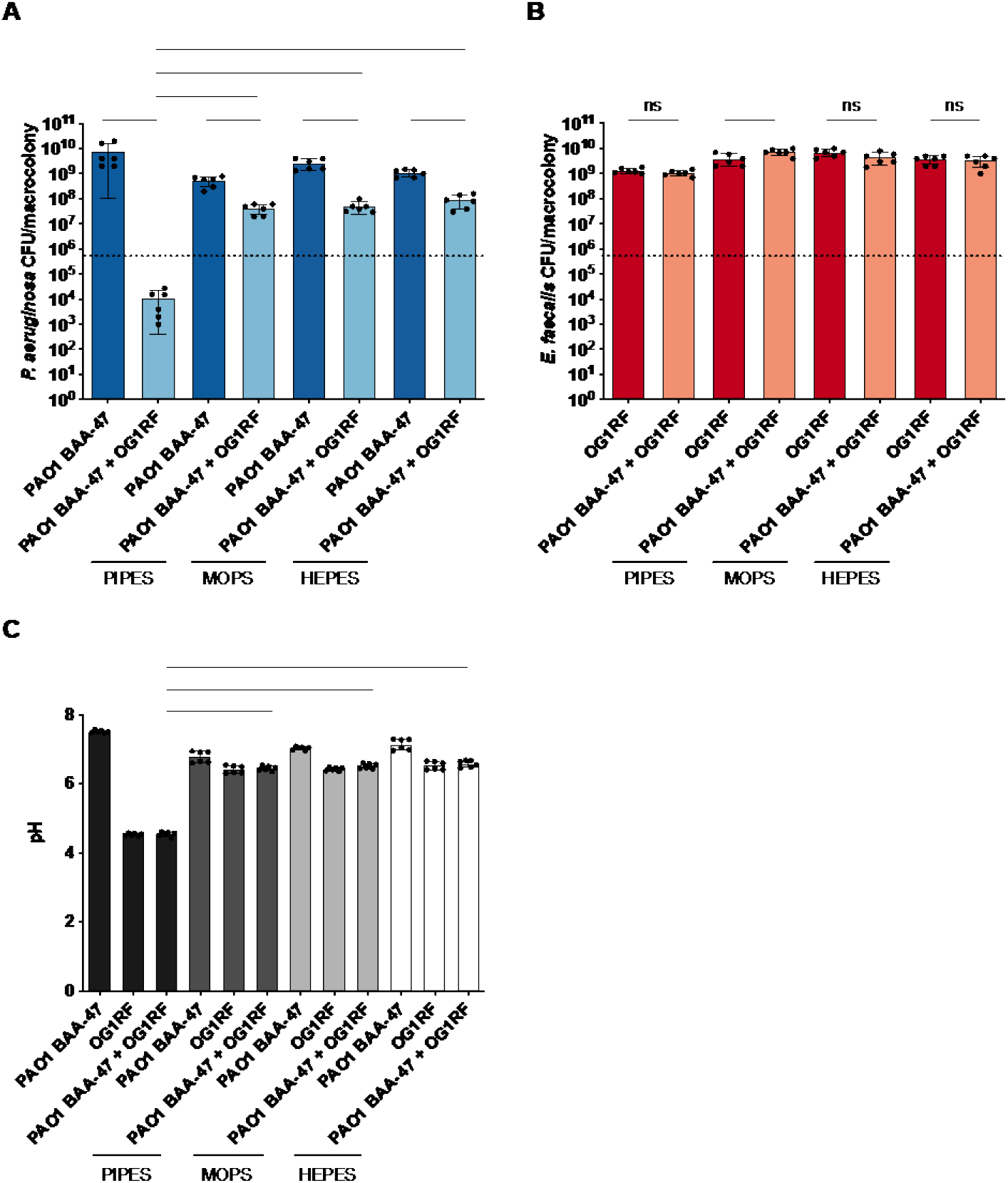
Alleviating pH acidification using buffered media partially rescues *P. aeruginosa* growth inhibition in iron-restricted conditions. Enumeration of **(A)** PAO1 BAA-47 and **(B)** OG1RF from single- and mixed-species macrocolonies grown for 24 h in media supplemented with 1 mM 22D and either 200 mM PIPES, MOPS or HEPES buffer. Bacterial species were mixed at a 1:1 ratio for mixed-species macrocolonies. Dotted lines represent inoculum of bacteria spotted. N = 2 with 3 technical replicates; error bars represent SD from the mean. Statistical analysis was performed using Mann-Whitney U test, *p < 0.05, **p < 0.01, ***p < 0.001, ****p < 0.0001. **(C)** The corresponding pH quantification of single- and mixed-species macrocolonies in **(A)** and **(B)**. Error bars represent SD from the mean. Statistical analysis was performed using Mann-Whitney U test, *p < 0.05, **p < 0.01, ***p < 0.001, ****p < 0.0001.

**Supplementary Table 1.**
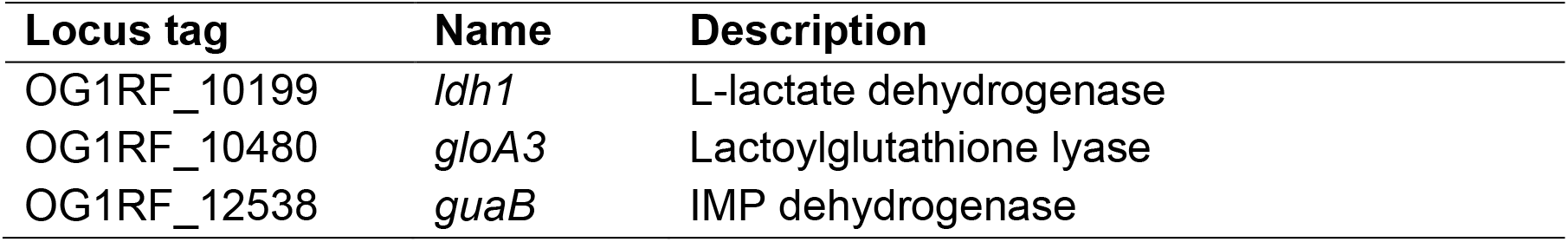
*E. faecalis* transposon mutants identified from transposon library screen with PADP6 in iron-restricted conditions.

| Locus tag | Name | Description |
| --- | --- | --- |
| OG1RF_10199 | *ldh1* | L-lactate dehydrogenase |
| OG1RF_10480 | *gloA3* | Lactoylglutathione lyase |
| OG1RF_12538 | *guaB* | IMP dehydrogenase |

**Supplementary Table 2.**
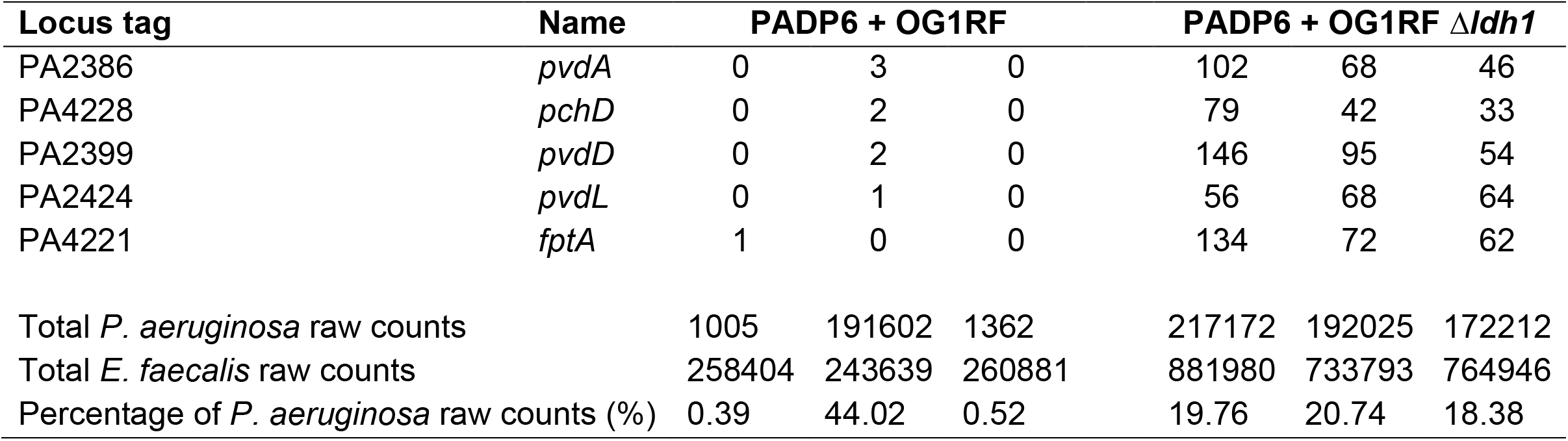
Selected *P. aeruginosa* raw counts of iron acquisition genes from mixed PADP6 and OG1RF macrocolonies and mixed PADP6 + OG1RF Δ*ldh1* macrocolonies in iron-restricted conditions.

**Supplementary Table 3.**
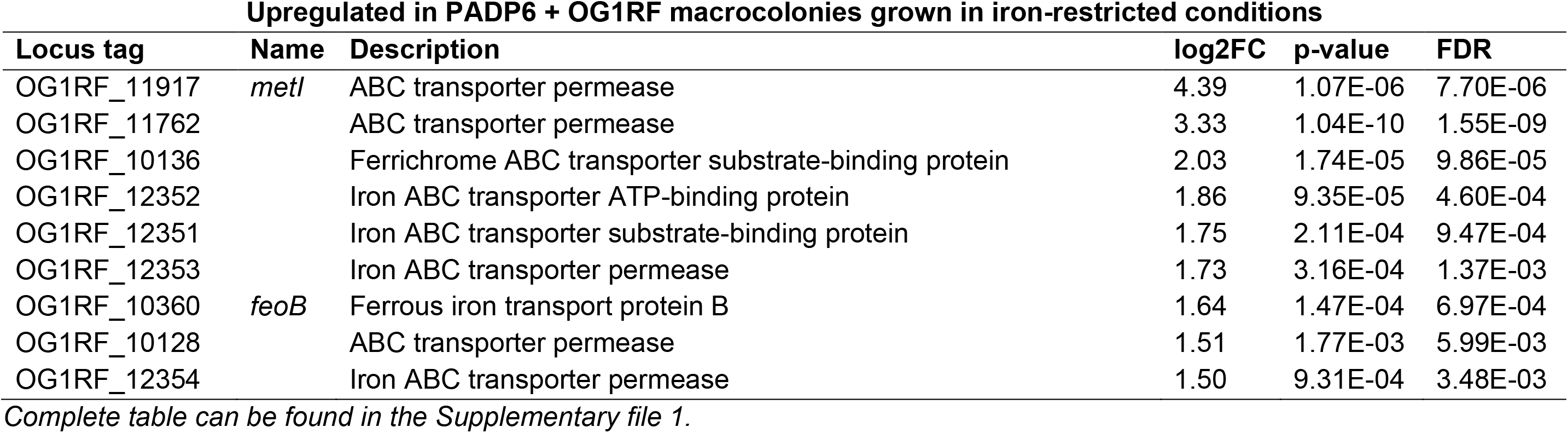
*E. faecalis* iron acquisition genes upregulated in PADP6 + OG1RF macrocolonies relative to PADP6 + OG1RF Δ*ldh1* macrocolonies grown in iron-restricted conditions.

**Supplementary Table 4.** Bacterial strains used in this study.

| Strain | Reference |
| --- | --- |
| *P. aeruginosa* |  |
| PAO1 WT | (100) |
| PADP6 | This study |
| PADP6-mCherry | This study |
| PAO1 BAA-47 | (101) |
| PR1 | (101) |
| PR2 | (101) |
| PR30 | (101) |
| PR80 | (101) |
| PR82PR101 | (101) |
| PR305 | (101) |
| PR325 | (101) |
| *E. faecalis* |  |
| OG1RF | (102) |
| OG1RF $\Delta ldh1$ | This study |
| OG1RF $\Delta ldh1::ldh1$ | This study |
| DS16 MG1505 ST40 | (103) |
| Merz 192 MG1519 ST40 | (103) |
| WH257 MG1422 ST9 | (103) |
| WH571 MH1424 ST9 | (103) |
| SF26630 MG1556 ST6 | (103) |
| V583MG899 ST6 | (103) |
| T13 MG1344 ST21 | (103) |
| SF24396 MG1544 ST21 | (103) |
| UAA1014 MG2962 ST23 | PRJNA88811 |
| B84847 MG3030 ST23 | (104) |
| SF21521 MG1542 ST28 | (103) |
| B56765 MG3031 ST28 | (104) |
| *E. coli* |  |
| *E. coli* pTNS1 | (93) |
| *E. coli* pUC18-miniTn7-P<sub>tac</sub>-mCherry | (105) |

**Supplementary Table 5.**
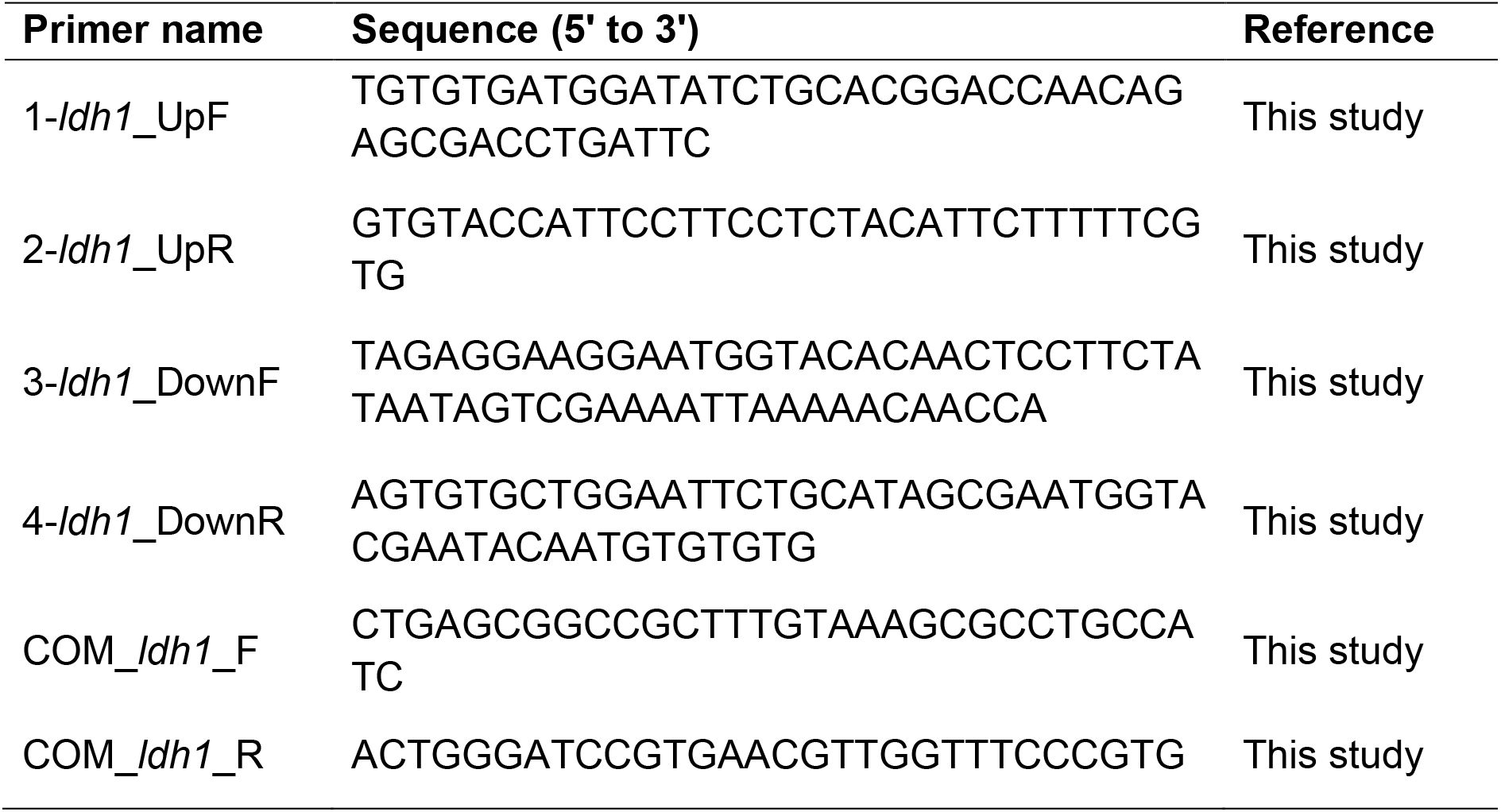
Primers used in this study.

